# Effect of Shufeng Xingbi Therapy on Th1/Th2 immune balance and intestinal flora in rats with allergic rhinitis

**DOI:** 10.1101/2025.03.26.645398

**Authors:** Shuiping Yan, Jian Zheng, Lizhen Huang, Yimin Zhou, Si Ai, Xiaoqiang Xie, Lihong Chen, Xiangli Zhuang, Minfeng Yu

## Abstract

**Objective:** To study the effect of Shufeng Xingbi Therapy (SFXBT) on Th1/Th2 immune balance and intestinal flora in allergic rhinitis (AR) rats.

**Methods:** 32 SD rats were randomly divided into control group, OVA group, antibiotic + SFXBT group and acetic acid + SFXBT group. Evaluate the AR behavioral score, observe the pathological changes of nasal mucosa with H&E, detect the composition and content of colonic contents with 16S rDNA, the levels of serum IgE, IL-4 and short-chain fatty acid (SCFAs) with ELISA, the mRNA expressions of STAT5, STAT6 and GATA3 in nasal mucosa with RT-qPCR, and the protein expressions of IL-4, STAT5, STAT6 and GATA3 in nasal mucosa with Western Blot.

**Results:** Compared with the OVA group, the AR behavioral score in the antibiotic + SFXBT group and acetic acid + SFXBT group decreased (*P* < 0.01), and the pathological changes of nasal mucosa were alleviated. At the door level, the relative abundance of *Firmicutes* in feces increased significantly, while the relative abundance of *Bacteroidetes* decreased significantly. At the genus level, The relative abundance of fecal *Lactobacillus, Romboutsia, Allobaculum* and *Dubosiella* increased significantly, the levels of serum IgE and IL-4 decreased (*P* < 0.05), the content of SCFAs increased significantly (*P* < 0.05), and the expression levels of STAT5, STAT6 and GATA3 mRNA and protein in nasal mucosa decreased significantly (*P* < 0.05).

**Conclusion:** Shufeng Xingbi Therapy can significantly improve the inflammatory symptoms of nasal mucosa in AR rats, and its mechanism may be closely related to regulating Th1/Th2 immune balance and intestinal flora.

## Introduction

The typical clinical symptoms of allergic rhinitis (AR) are paroxysmal sneezing, watery nasal discharge, nasal itching and nasal congestion, which is a non-infectious chronic inflammatory disease of nasal mucosa mainly mediated by IgE after the body is exposed to allergens [1]. The prevalence of AR in the world is over 10%, and it has risen rapidly in the past decades, which has seriously affected the quality of life of patients and increased their economic burden [2].

At present, it is believed that the pathogenesis of AR is mainly caused by Th1/Th2 immune imbalance. The treatment of AR in modern medicine includes drug therapy, immunotherapy and surgical treatment, among which glucocorticoid, antihistamine, leukotriene receptor antagonist and decongestant are commonly used. These drugs can improve the symptoms of AR in children in a short time, but to some extent, they will bring local or even systemic adverse reactions and side effects to sick children [3].

“hygiene hypothesis” links early exposure to environment and microorganisms with the prevalence of atopic allergy and asthma. Exposure to environment and microorganisms can affect the growing immune system and protect the subsequent immune-mediated diseases [4]. Unbalanced intestinal flora can lead to abnormal respiratory allergic reactions. In the lung, SCFAs, as a signal molecule of antigen presenting cells, participates in regulating allergic reactions and lung inflammation [5]. The ideal strategy to prevent and treat AR is to inhibit the excessive activation of local inflammatory reaction in nasal mucosa, and its mechanism needs to be further explored.

Traditional Chinese Medicine (TCM) has obvious advantages in treating children’s AR. In recent years, it has achieved good effects in treating AR and regulating the immune function of the body. Professor Zheng Jian concluded decades of clinical experience that the etiology and pathogenesis of TCM is pathogenic wind invading the lungs, and advocated the method of Shufeng Xingbi Therapy. Therefore, it was given Shufeng Xingbi recipe for oral administration and Xingbi gel nasal drops for external use, which was proved to be effective clinically. In this paper, the effects of Shufeng Xingbi Therapy on Th1/Th2 balance and intestinal flora in AR rats were studied, and the mechanism of Shufeng Xingbi recipe for oral administration combined with nasal drops for external use in treating AR was discussed, so as to provide experimental basis for Shufeng Xingbi Therapy in treating AR.

## Materials and Methods

### Animals and treatment

32 clean healthy male SD rats, 6 weeks old and weighing 200 - 250 g, were ordered from Shanghai slack Experimental Animal Co., Ltd. with the license number of SCXK (Shanghai) 2012 - 0011. The experimental rats are kept in the Experimental Animal Center of Fujian University of Traditional Chinese Medicine [license number: SYXK (Fujian) 2020 - 0003], which meets the basic conditions of animal feeding. Feeding conditions: temperature 22 °C, humidity 50% ∼ 70%, simulating standard day and night system (12h light +12h night), giving free diet and drinking water. This experiment has been approved by the Ethics Committee of Fujian University of Traditional Chinese Medicine (approval number: 2021022). During the experiment, the experimental animals were disposed according to the Guiding Opinions on Treating Experimental Animals Well issued by the Ministry of Science and Technology of the People’s Republic of China.

### Ovalbumin (OVA) - induced AR model

32 rats were randomly divided into control group, OVA group, antibiotic + SFXBT group and acetic acid + SFXBT group, with 8 rats in each group. According to reference [6], AR rat model was established by OVA (A5503, Sigma Company, USA) + aluminum hydroxide sensitization (239186, Sigma Company, USA) and OVA stimulation. ① Sensitization: rats in OVA group, antibiotic + SFXBT group and acetic acid + SFXBT group were injected with 1mL of suspension containing 0.5 mg/mL OVA + 30 mg/mL aluminum hydroxide once every other day for 7 times, and the control group was injected with the same amount of 0.9% sodium chloride solution; ② Stimulation: On the 4th day after intraperitoneal injection sensitization, 50 μL of 2% OVA solution was dripped into each nasal cavity of rats once every other day for 5 times, while the control group was dripped with the same amount of 0.9% sodium chloride solution.

### Drug intervention

According to references [7-8], the flora was exhausted and supplemented with acetic acid. From the first day after intraperitoneal injection sensitization, rats in antibiotic + SFXBT group and acetic acid + SFXBT group were given compound antibiotic solution and compound antibiotic solution added acetic acid instead of pure water for daily drinking separately. The compound antibiotics contained neomycin (C3090, Apexbio, USA) 1 g/L, metronidazole (B1795, Apexbio, USA) 1 g/L, vancomycin (B1976, Apexbio, USA) 500 mg/L and polymyxin B (C6417, Apexbio, USA) 200 mg/L, which were dissolved in 1 L of purified water, while acetic acid (S5636, Sigma Company, USA) was dissolved in the compound antibiotic solution according to the concentration of 150 mM. Rats in antibiotic + SFXBT group and acetic acid + SFXBT group were both given 1 mL/100 g of Shufeng Xingbi recipe solution by stomach every day, and 50 μL/nostril of Xingbi gel nasal drops were given once a day for 14 days. Rats in control group and OVA group were given the same amount of 0.9% sodium chloride solution by stomach and drank pure water every day. Animal modeling and intervention process are shown in Figure 1.

**Figure 1.**
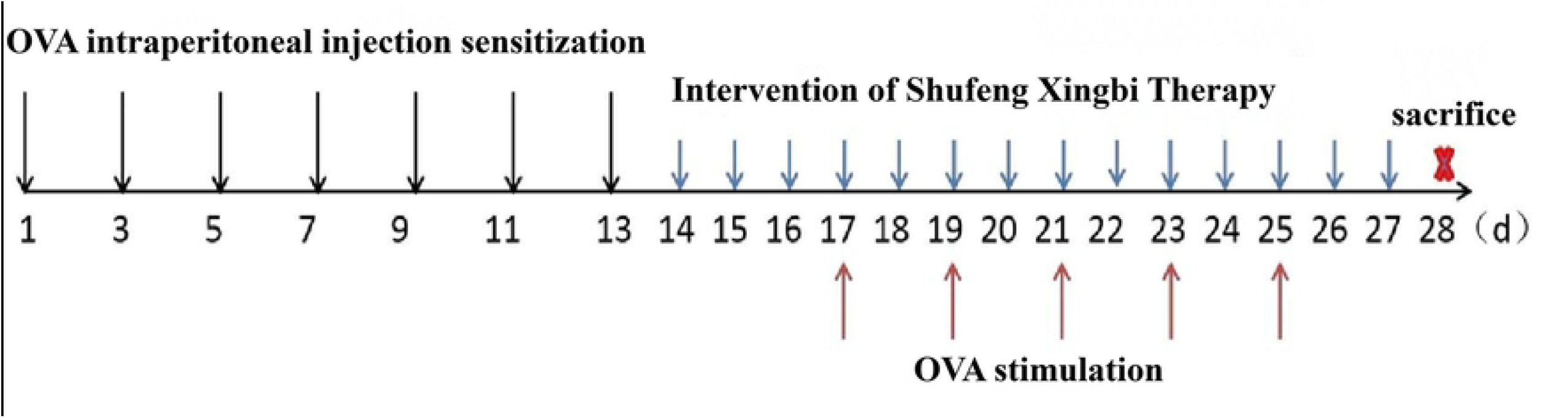
Animal modeling and intervention process.

Shufeng Xingbi recipe is composed of ephedra sinica 9 g, bitter almond 9 g, Pogostemon cablin 6 g, Ligusticum chuanxiong 5 g, loofah sponge 12 g, dandelion 15 g, Flos Magnoliae 9 g, raw licorice 5 g. It is made into a medicinal liquid with a crude drug concentration of 0.63 g/mL by filtration, and the content and quality control of ephedrine hydrochloride and amygdalin were carried out by high performance liquid chromatography, which was completed by the preparation room of Shenzhen Traditional Chinese Medicine Hospital. Xingbi gel nasal drops is a hospital preparation of People’s Hospital affiliated to Fujian University of Traditional Chinese Medicine (Z20110006), which is composed of Xu Changqing, Periostracum Cicadae, artificial bezoar and borneol. All of them are provided by the preparation room of the People’s Hospital affiliated to Fujian University of Traditional Chinese Medicine.

### Behavioral assessment of nasal symptoms

After each nasal drip of OVA, the times of sneezing, nasal scratching and nasal discharge of rats within 30 min were observed and recorded, and the total score was calculated by superposition method. The total score ≥ 5 indicates the success of modeling [9], and the scoring method was 1 point: sneezing 3-9 times, nasal itching/nasal scratching 2-3 times, and nasal discharge to nostrils; 2 points: sneezing 10-14 times, itchy nose/scratching nose 4-5 times, runny nose exceeding the front nostril; 3 points: sneezing is greater than or equal to 15 times, nasal itching/scratching is greater than 5 times, and the nose is full of tears.

### Hematoxylin and eosin (H&E) staining assay

The nasal tissue was fixed with 4% paraformaldehyde (PFA), embedded in paraffin, and cut into 5 μm thick slices. The sections were stained with hematoxylin and eosin (H&E; G1140 G110; Beijing suolaibao technology co., ltd, Beijing, China), and examined under the optical microscope (Leica company of Germany).

### NEB Next®Ultra™DNA Library preparation and illumina MiSeq sequencing

Extraction of genomic DNA: CTAB/SDS method was used to extract total genomic DNA from samples. The concentration and purity of DNA were monitored on 1% agarose gel. According to the concentration, the DNA was diluted to 1 ng/μL with sterile water. Amplicon primer: 16s v3-v4: 341f-806r, 18S V9: 1380F-1510R, ITS1: ITS1F-ITS2R. The 16S /18S rRNA gene was amplified with specific primers with bar codes. All PCR reactions were carried out in 30 μL of reactants with 15 μL of Phusion High-Fidelity PCR Master Mix (New England Biological Laboratory), with 0.2 μM of forward and reverse primers and about 10 ng of template DNA. Thermal cycling includes initial denaturation at 98 °C for 1 min; Denaturing at 98 °C for 10 s, annealing at 50 °C for 30 s, and extending at 72 °C for 60 s for 30 cycles; Finally, 72 °C for 5 min.Quantitative and qualitative analysis of PCR products 1× loading buffer (containing SYB green) with the same volume was mixed with PCR products, and electrophoresis detection was carried out on 2% agarose gel. Select the samples whose main band color is between 400 and 450 BP for further experiments. Mixing and purification of PCR products The PCR products were mixed in equal density ratio. Then, the mixed PCR products were purified by using AxyPrepDNA gel extraction kit (AXYGEN Company). According to the manufacturer’s suggestion, NEB NextTulatmDNA Library PrepKit was used to prepare sequencing library for Illumina, and the index code was added. The quality of the library was evaluated by using Qubit@ 2.0 Fluorometer and Agilent Bioanalyzer 2100 system. Finally, the library was sequenced on Illumina NovaSeq 600 platform, and a 250bp double-ended reading was generated.

### Enzyme-Linked Immunosorbent Serological Assay (ELISA)

Serum levels of IgE, IL-4, SCFAs were measured by using ELISA kits (Shanghai enzyme-linked organism, China) according to the manufacturer’s instructions.

### RNA extraction and real-time quantitative polymerase chain reaction (RT-qPCR)

Total RNA was extracted with Trizol (15596026, American Life company), and the extracted RNA was reverse transcribed according to the instructions of the kit to synthesize cDNA (R211-02, Q711-02, Nanjing nuoweizan biology, China). The mRNA expressions of STAT5, STAT6 and GATA3 were detected by RT-qPCR using the cDNA as a template. Reaction conditions: 95 °C, 30 s; 95 °C 、 10 s; 60 °C, 30 s, 40 cycles in total. Using β-actin as internal reference, the relative expression of related gene mRNA was calculated by 2^-ΔΔ^ CT method. The sequences of various primers are shown in Table 1.

**Table 1.**
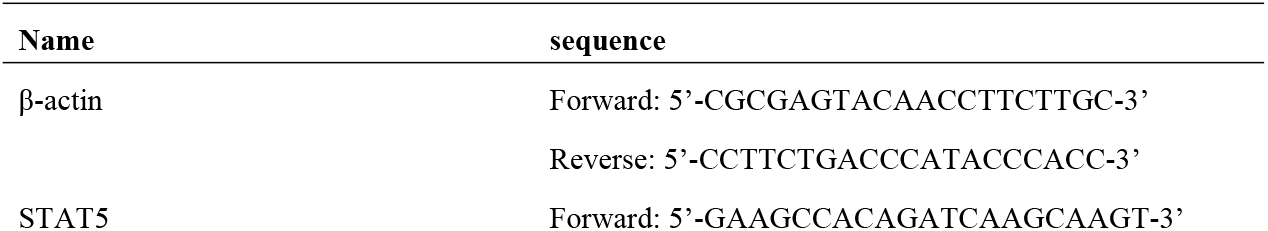
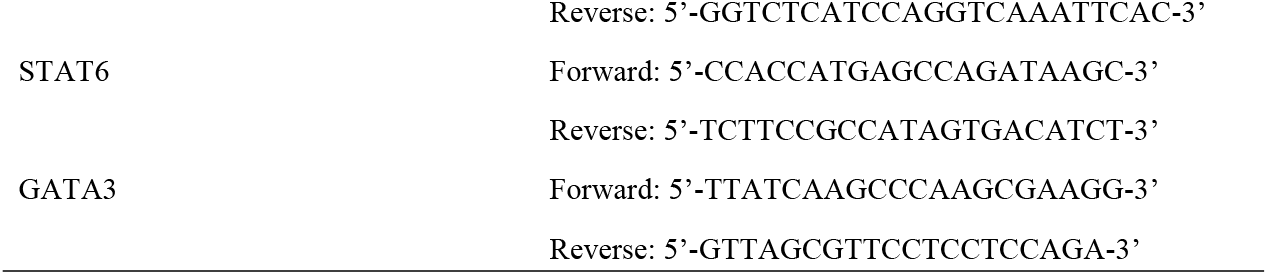
Primer sequences.

### Western blotting assay

Take nasal mucosa tissue and add a proper amount of lysis buffer, homogenize and lyse it on ice for 30 min, then centrifuge at 4 °C and 12000 rpm for 30 min, then take albumin, and measure the protein concentration by BCA method. 30 μg of protein was loaded, separated by SDS-PAGE electrophoresis, transferred to solid support PVDF membrane, sealed with sealing liquid, incubated with the corresponding primary antibodies: anti-rabbit STAT6 (1:1000, 51073; Proteintech, China), anti-mouse IL-4 (1:1000, 66142; Proteintech, China), anti-mouse STAT5 (1:1000, ab230670; abcam, USA), anti-rabbit GATA2 + GATA3 (1:1000, ab182747; abcam, USA), and anti-mouse β-actin (1:20000, 66009; Proteintech, China), The PVDF membranes were washed, and secondary antibodies were applied in ratio of 1:2000 for 1 h at room temperature, and imaged by ECL chemiluminescence. Using β-actin as internal reference, the optical density values of hybrid protein bands were compared and analyzed by analysis software.

### Statistical analysis

Statistical analyses were performed using One-way ANOVA and Student’s t-test. Data were presented as mean values ± standard error of mean (SEM) of at least three independent experiments. *P* ≤ 0.05 were considered as significant.

## Results

### Shufeng Xingbi Therapy reduces nasal symptoms in OVA - induced AR rat

After modeling, as shown in Figure 2, the behavioral scores of rats in OVA group, antibiotic + SFXBT group and acetic acid + SFXBT group all exceeded 5 points, which was significantly higher than that in control group (*P* < 0.01). After the intervention of TCM, the rats were scored again within 30 minutes. Compared with the OVA group, the behavioral scores of the rats in the antibiotic + SFXBT group and acetic acid + SFXBT group were significantly lower (*P* < 0.01), and both were significantly lower than those before the intervention (*P* < 0.01), and there was no significant difference compared with the control group (*P* > 0.05). These results implied that Shufeng Xingbi Therapy alleviated nasal symptoms.

**Figure 2.**
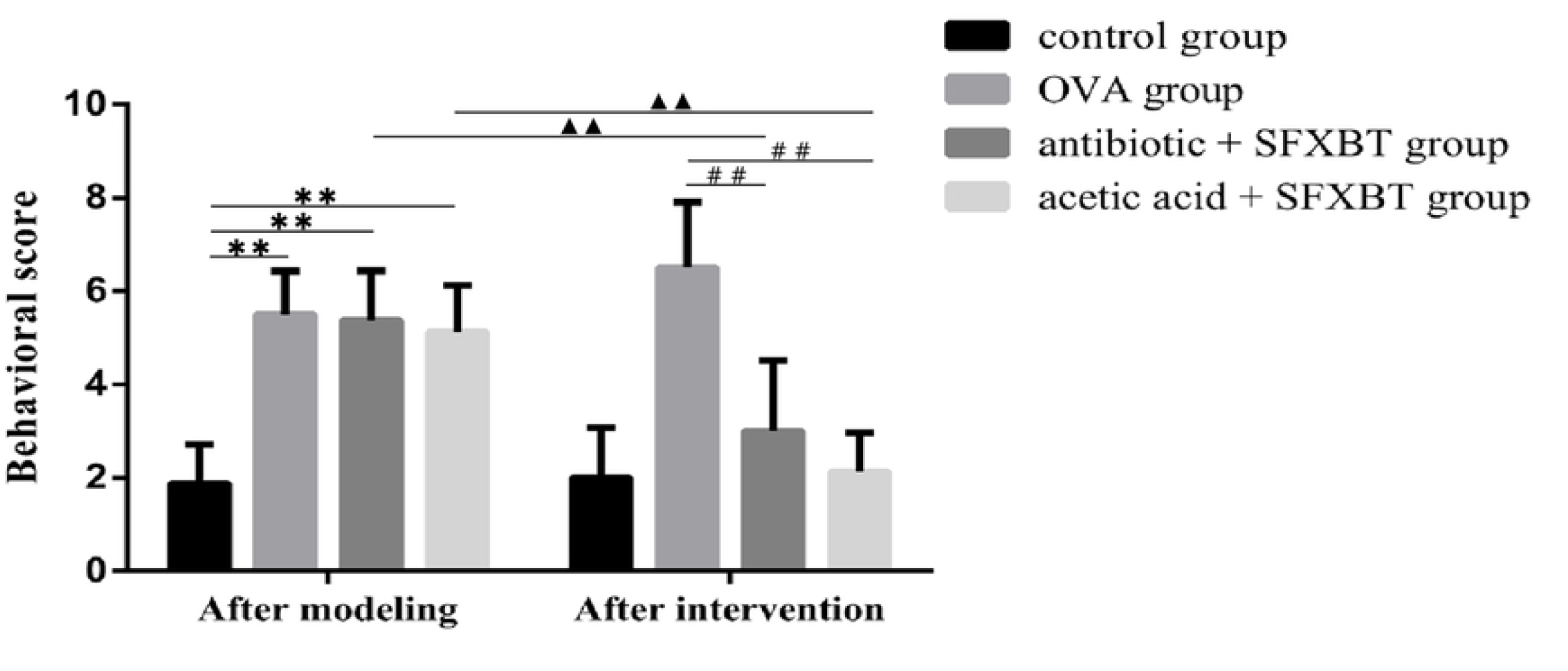
The behavioral scores of rats. \*\**P* < 0.01,^▲▲^*P* < 0.01,^##^*P* < 0.01. Data were shown as the mean ± SD, n = 3 in each group, and all by analysis of variance.

### Shufeng Xingbi Therapy improves the pathological changes of nasal mucosa in OVA-induced AR rat

As shown in Figure 3, the epithelial structure of nasal mucosa in the control group was smooth and complete, arranged neatly, and the cilia were continuous and complete, without obvious vasodilation. In the OVA group, most nasal cilia fell off, the mucosal structure was incomplete, the epithelial cells were arranged in disorder, a large number of inflammatory cells were infiltrated, and the glands and blood vessels around submucosa expanded and proliferated obviously. After the treatment of Shufeng Xingbi Therapy, the edema of nasal mucosa in antibiotic + SFXBT group and acetic acid + SFXBT group was obviously reduced, the epithelial cell structure was relatively complete, the cilia were relatively continuous and complete, there was no obvious congestion and expansion of blood vessels, and a small amount of inflammatory cells could still be seen, but the pathological changes were obviously improved compared with the OVA group. These results implied that Shufeng Xingbi Therapy alleviated the pathological changes of nasal mucosa.

**Figure 3.**
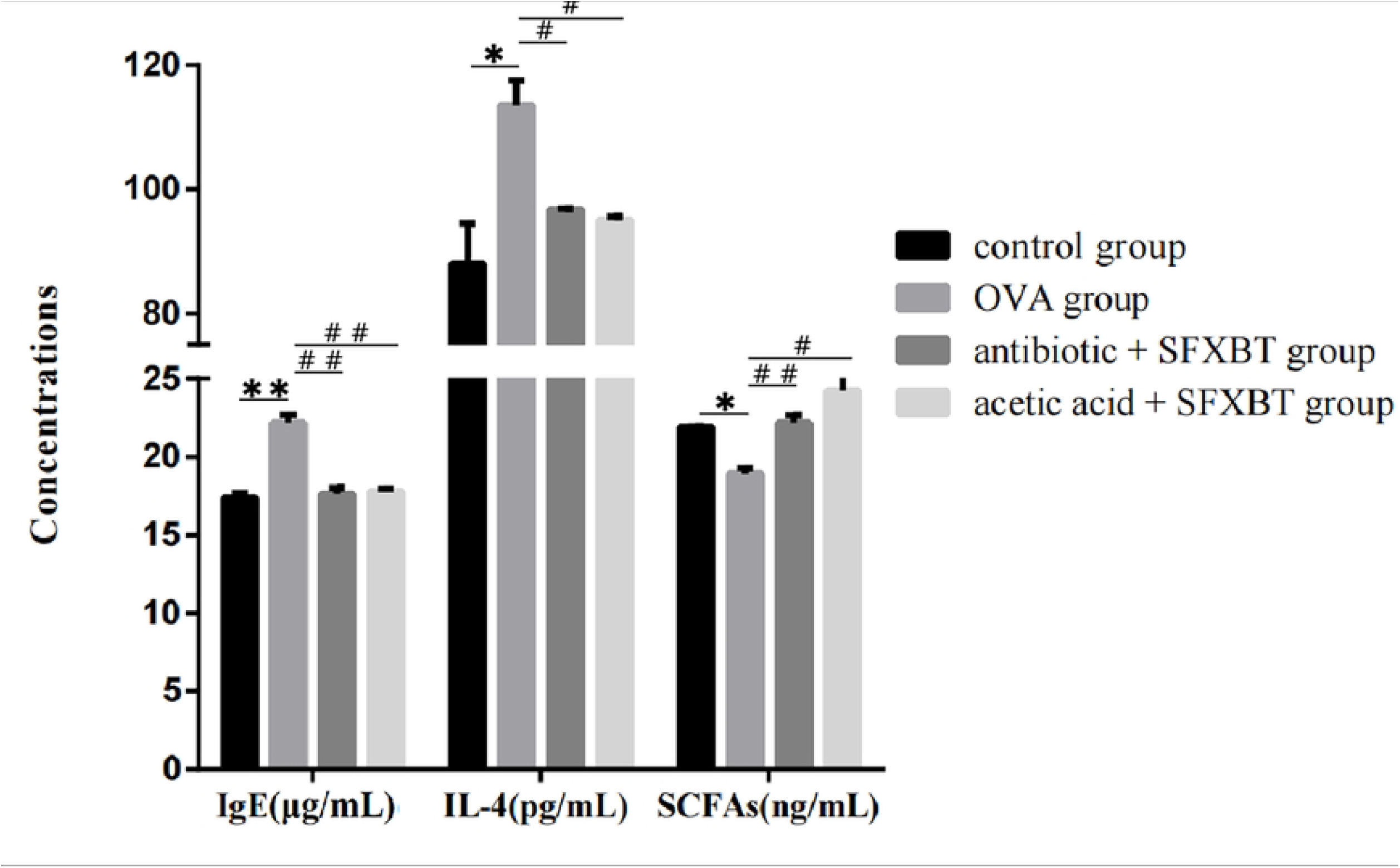
Histopathological changes in hematoxylin and eosin (H&E) (×200). A.control group, B.OVA group, C.antibiotic + SFXBT group, D.acetic acid + SFXBT group.

### Shufeng Xingbi Therapy improves intestinal flora composition and SCFAs content in serum in OVA-induced AR rat

As shown in Figure 4, at the door level, compared with the control group, the relative abundance of *Firmicutes* in the feces of rats in the OVA group decreased significantly, while the relative abundance of *Bacteroidetes* increased significantly. After the intervention of Shufeng Xingbi Therapy, compared with the OVA group, the relative abundance of *Firmicutes* in the feces of rats in the antibiotic + SFXBT group and acetic acid + SFXBT group increased significantly, while the relative abundance of *Bacteroidetes* decreased significantly. At the genus level, the relative abundance of *Lactobacillus* and *Romboutsia* in the feces of rats in the OVA group decreased significantly, while the relative abundance of *Allobaculum* and *Dubosiella* in the OVA group also decreased to varying degrees. The relative abundance of *Lactobacillus, Romboutsia, Allobaculum* and *Dubosiella* can be increased after the intervention of Shufeng Xingbi Therapy. As shown in Figure 5, the content of SCFAs in serum was detected by ELISA. Compared with the control group, the content of SCFAs in serum in OVA group decreased (*P* < 0.05), After the intervention of Shufeng Xingbi Therapy, compared with the OVA rats, the content of SCFAs was significantly increased (antibiotic + SFXBT group *P* < 0.01, acetic acid + SFXBT group *P* < 0.05). These results implied that Shufeng Xingbi Therapy improves the intestinal flora composition and SCFAs content in serum.

**Figure 4.**
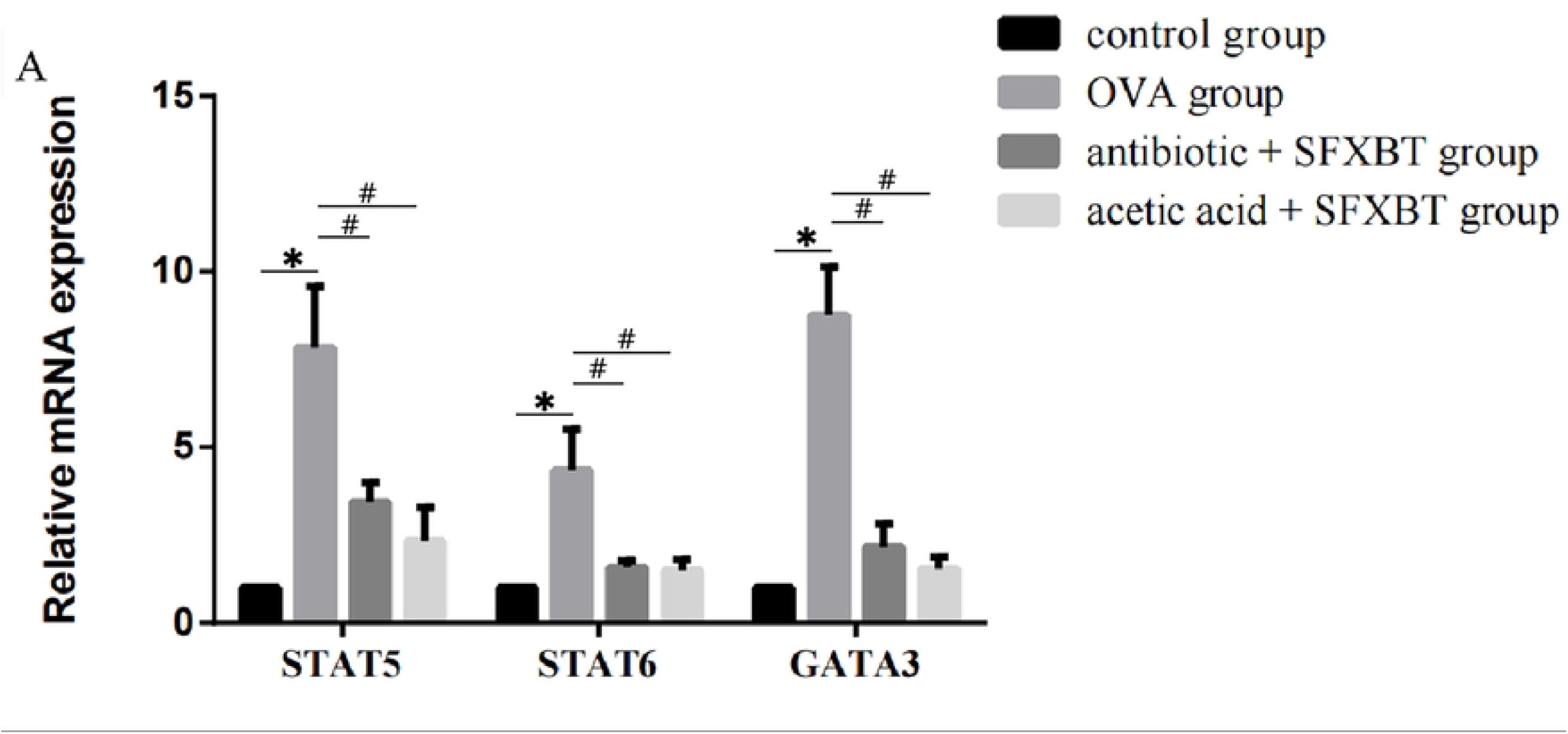
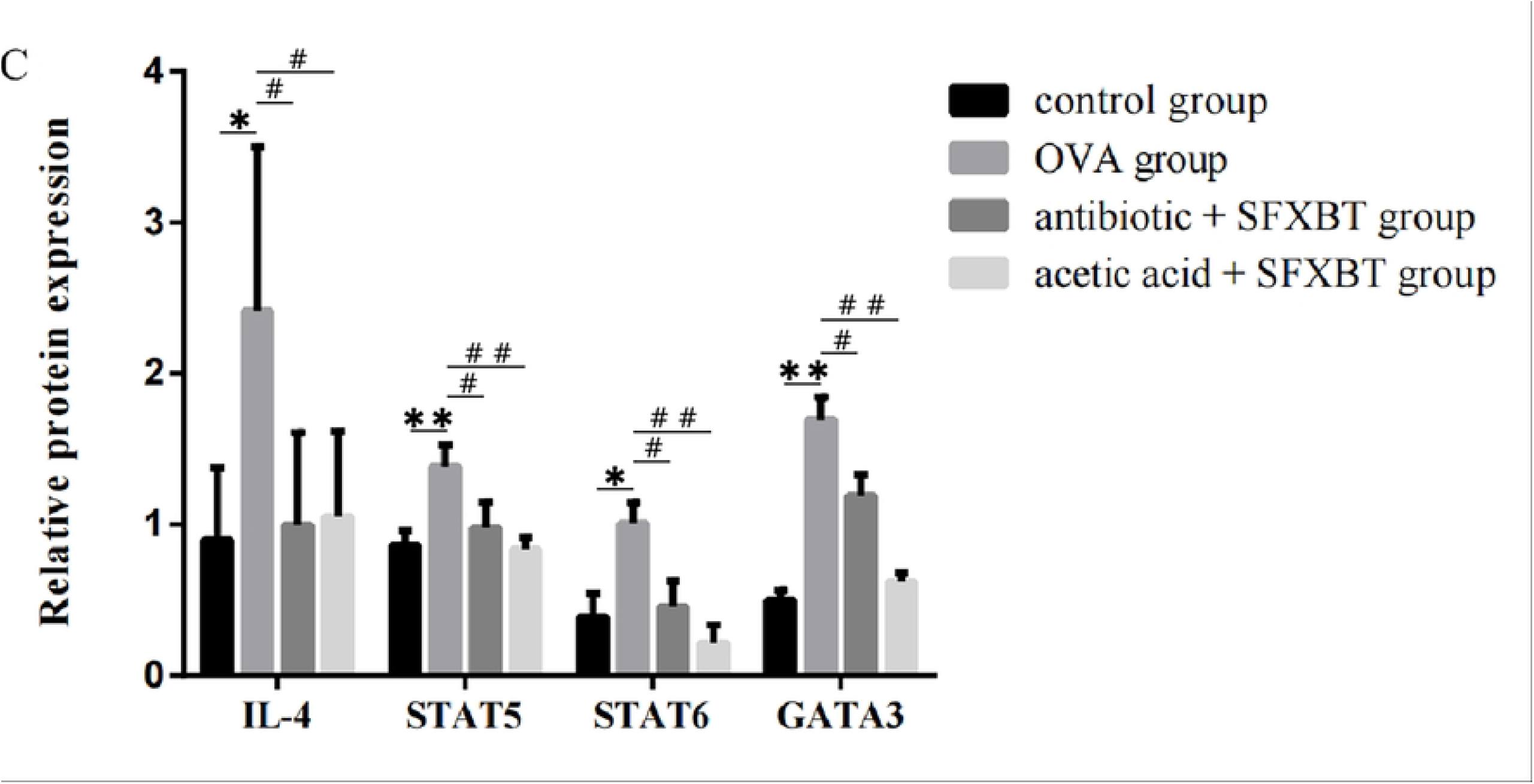
Intestinal flora composition. The relative abundance of flora at the door (left) and genus (right) levels in rat colon.

**Figure 5.**
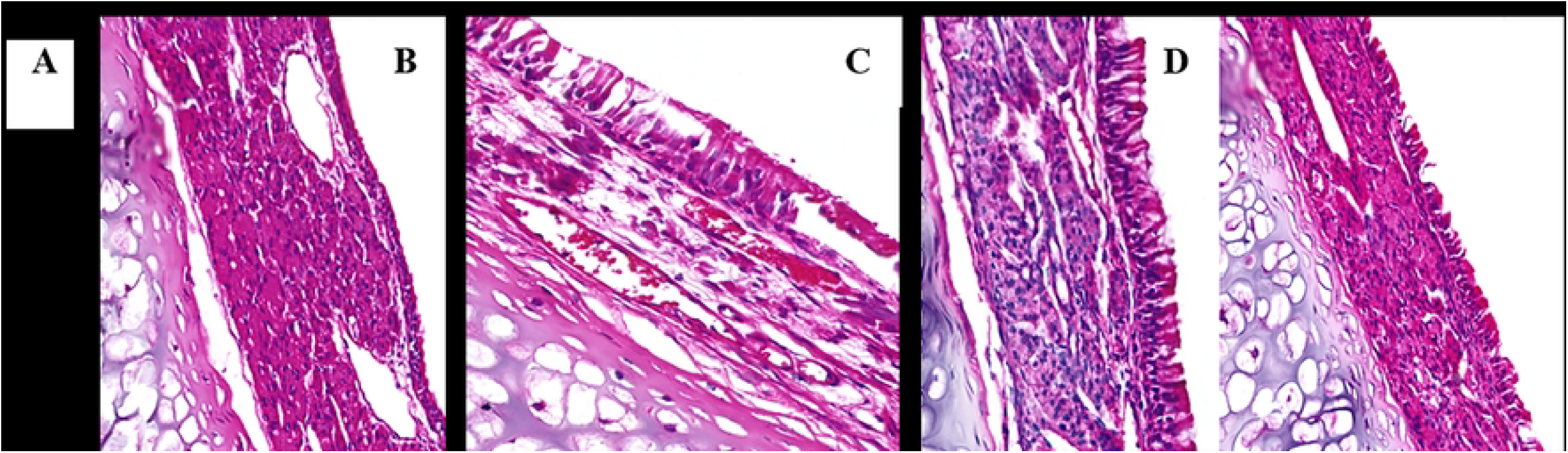
Serum levels of IL-4, IgE and SCFAs. \**P* < 0.05, \*\**P* < 0.01,^#^*P* < 0.05,^##^*P* < 0.01. Data were shown as the mean ± SD, n = 3 in each group, and all by analysis of variance.

### Shufeng Xingbi Therapy suppresses inflammatory cytokine production in AR rat

Inflammatory response in OVA - induced AR rat was assessed by measuring serum levels of IL-4, IgE by using ELISA. As shown in Figure 5, compared with the control group, the contents of serum IgE and IL-4 in the OVA group increased significantly (IgE *P* < 0.01, IL-4 *P* < 0.05). After the intervention of Shufeng Xingbi Therapy, compared with the OVA rats, the levels of serum IgE and IL-4 in the antibiotic + SFXBT group and acetic acid + SFXBT group were significantly decreased (IgE *P* < 0.01, IL-4 *P* < 0.05).

### Shufeng Xingbi Therapy regulates Th1/Th2 immune balance in AR rat

In order to clarify the mechanism of regulating Th1/Th2 immune balance by Shufeng Xingbi Therapy, we analyzed the expression of Th2 cell-related inflammatory factors by western blot and RT-qPCR. As shown in Figure 6, the mRNA and protein levels of STAT5, STAT6 and GATA3 in the OVA group were significantly higher than those in the control group (*P* < 0.05). After the intervention of Shufeng Xingbi Therapy, the mRNA and protein expressions of STAT5, STAT6 and GATA3 in the antibiotic + SFXBT group and acetic acid + SFXBT group were significantly decreased (*P* < 0.05), and the expression of IL-4 protein was also significantly decreased (*P* < 0.05). The results show that Shufeng Xingbi Therapy can improve AR by regulating Th1/Th2.

**Figure 6.**
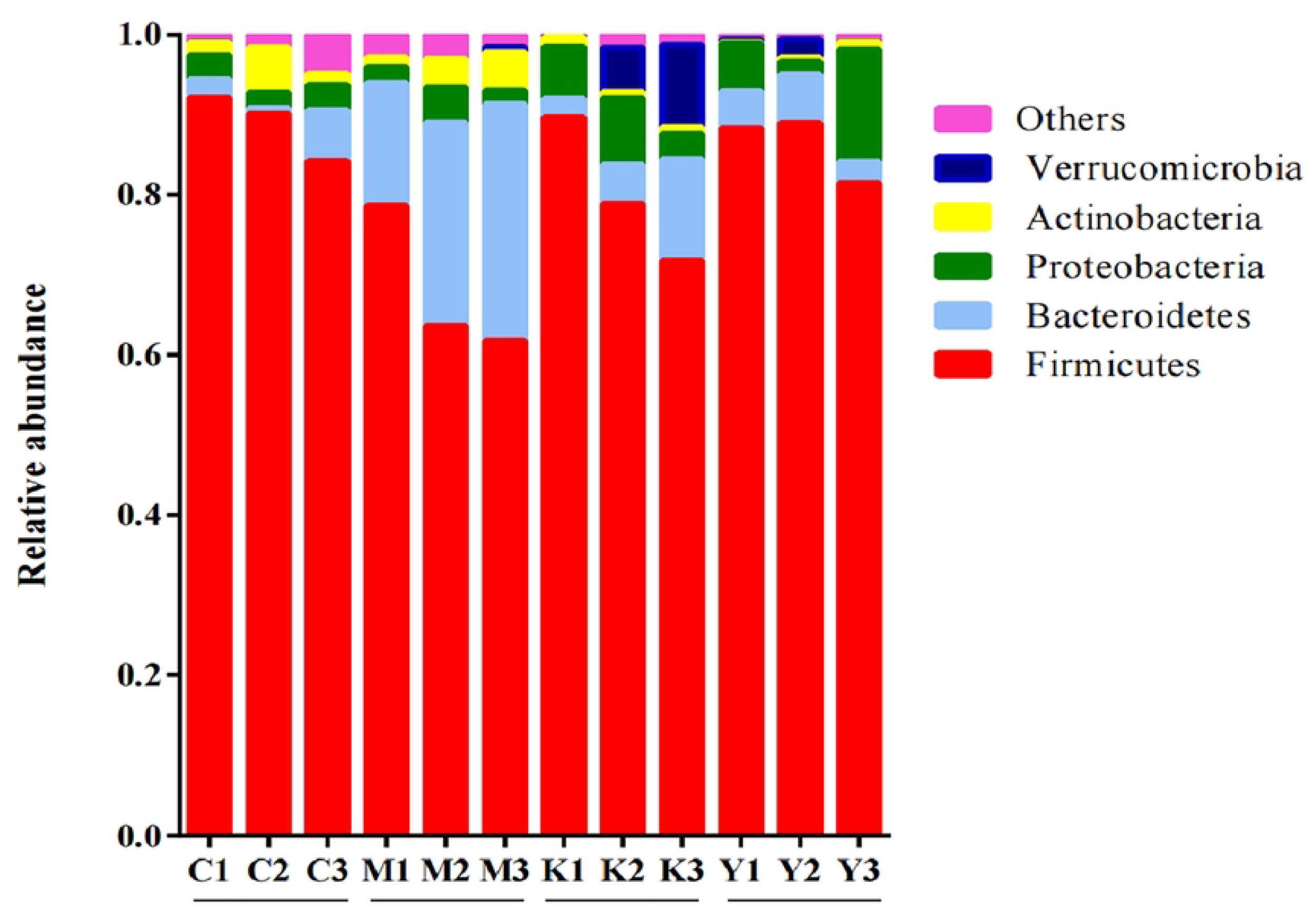
The mRNA and protein levels of STAT5, STAT6 and GATA3. (A)Expression of STAT5, STAT6 and GATA3 mRNA in nasal mucosa. (B-C)Expression of IL-4, STAT5, STAT6 and GATA3 proteins in nasal mucosa.\**P* < 0.05, \*\**P* < 0.01,^#^ *P* < 0.05,^# #^ *P* < 0.01. Data were shown as the mean ± SD, n = 3 in each group, and all by analysis of variance.

## Discussion

AR belongs to type I allergic reaction, and its pathogenesis is complicated. At present, a large number of studies have confirmed that the pathogenesis of AR is related to the dominant response of Th2 cells caused by Th1/Th2 immune imbalance. Th2 cells can produce inflammatory factors such as IL-4, IL-10, IL-13 and IL-5, among which IL-4 and IL-13 can promote the synthesis of IgE. IL-4 mediated by Th2 cytokines plays an important role in the pathogenesis of AR and is the key guiding signal of its polarization [10]. Signal transducers and activators of transcription (STAT) play an important role in regulating the dynamic balance, differentiation and cell function of immune cells. STAT6 can regulate allergic reactions, which are mainly activated by IL-4 and IL-13. Besides promoting the differentiation of Th2 cells, STAT6 also plays a vital role in promoting the development, differentiation and class switching of B cells [11]. IL-4 induces Th2 cell differentiation through the transcription factor STAT6, and STAT6 induces the expression of GATA3, which is the central transcription factor necessary for Th2 differentiation and cytokine production. GATA3 and STAT5 activated by IL-2, IL-7 or TSLP signal combine to form a positive feedback loop for amplifying and strengthening Th2 gene expression programs. Many of these changes are related to the epigenetic modification of Th2 cytokine sites, mainly caused by GATA3, and the up-regulation of IL-2α and IL-4α receptor chains, thus making IL-2-STAT5 and IL-4-STAT6-GATA3 signaling pathways persist [12,13].

The hypothesis of intestinal microflora holds that AR occurs because a series of external factors change the structure of gastrointestinal flora, which leads to the impairment of the regulation of intestinal flora on host immune tolerance [14]. Through the common mucosal immune system, changes in the composition and function of the flora of the digestive tract can affect the respiratory system, while through immune regulation, the disorder of the flora of the respiratory tract can also affect the digestive tract. This interaction between the intestine and the lungs is called the “lung-intestine axis”. The two most important organ systems of the human body that communicate with the outside world are the digestive tract and the respiratory tract, and the relationship between the lung and the intestine has gradually attracted academic attention and formed the “intestine-lung axis” theory. In the lung, SCFAs, as a signal molecule of antigen presenting cells, is involved in regulating allergic reaction and lung inflammation [4]. Healthy intestinal microecology can prevent allergic reaction, mainly through the production of SCFAs or by acting on type3 intrinsic lymphocytes (ILCs) to produce IL-22, which can induce Paneth cell to produce antibacterial peptides (AMP) and goblet cells to produce mucus, thus enhancing the barrier function [15], and SCFAs can also inhibit immune inflammation through G protein-coupled receptors (GPCRS) [16-17], probiotics have a certain effect on the treatment of AR. At present, probiotics can not be recommended as an independent treatment for AR, but as a new potential treatment, people think that it can be used as an adjuvant treatment for AR [18]. Intestinal flora can regulate the immune response of respiratory tract and intestine through different regulatory pathways.

Shufeng Xingbi Therapy is a method of oral and external treatment with Shufeng Xingbi recipe combined with Xingbi gel nasal drops, which is the clinical experience of famous Chinese medicine practitioners in Fujian Province in treating AR. Through animal experiments, it is found that Shufeng Xingbi Therapy can reduce the expression of Fyn and SCF, up-regulate STAT5, antagonize IgE synthesis, correct Th1/Th2 immune imbalance and reduce the immune response of AR fibroblasts. Through clinical research, it is found that Shufeng Xingbi Therapy can inhibit the release of inflammatory mediators by down-regulating the levels of serum immune inflammatory factors such as IL-13, IgE, FcεRI, TSLP and IL-4, on the one hand, it can increase the level of INF-γ and regulate the differentiation of Th cells, on the other hand, it can promote the secretion of Th1 cytokines with anti-inflammatory effect by inhibiting IL-33/ST2 signaling pathway, so as to rebalance Th1 and Th2 responses to improve the symptoms of children with AR. After 4 weeks of treatment, the total effective rate is 96.07%.

The results of this experiment showed that AR rats induced by OVA+ aluminum hydroxide sensitization and OVA stimulation showed obvious inflammatory symptoms of nasal mucosa, a large number of inflammatory cells infiltrated in nasal mucosa, glands and blood vessels around submucosa were congested and edematous, fecal flora was disordered, serum IgE and IL-4 contents were increased, SCFAs expression was decreased, and the expression levels of STAT5, STAT6, GATA3 mRNA and IL-4, STAT5, STAT6 and GATA3 proteins in nasal mucosa were significantly increased. It is suggested that the expression levels of IL-4, STAT5, STAT6, GATA3 and intestinal flora play a key role in the occurrence and development of AR diseases, and Th2 immune response is dominant in Th1/Th2 imbalance. After the exhaustion of antibiotic flora, the intervention of Shufeng Xingbi Therapy and supplementing with acetic acid can effectively improve the nasal inflammatory symptoms of AR rats, obviously improve the infiltration of inflammatory cells in nasal mucosa, the proliferation and expansion of glands and blood vessels around submucosa, adjust the composition ratio of *Firmicutes* and *Bacteroidetes* at the door level, and increase the relative abundance of *Lactobacillus* at the genus level. The levels of serum IgE and IL-4 decreased significantly, and the expression of SCFAs increased. At the same time, the expression levels of STAT5, STAT6, GATA3 mRNA and IL-4, STAT5, STAT6 and GATA3 protein in nasal mucosa were decreased, so that the Th1/Th2 immune balance was transformed to Th1 immune response trend, and the intestinal flora structure was adjusted, thus regulating the immune imbalance of AR rats and improving symptoms.

## Conclusion

To sum up, the method of Shufeng Xingbi Therapy can significantly improve the inflammatory symptoms of nasal mucosa in AR rats, and its mechanism may be closely related to the regulation of Th1/Th2 immune balance and intestinal flora, which provides an important experimental basis for the clinical application of the method of Shufeng Xingbi Therapy to treat AR.

## Supporting information

**S1 Raw images. (PDF)**

## Declarations

Thanks to all those who have offered their help and support.

## Consent for publication

Written informed consent for publication was obtained from all participants.

## Data Availability Statement

All relevant data are within the manuscript and its Supporting Information files.

## Author Contributions

**Conceptualization:** Shuiping Yan, Jian Zheng, Xiangli Zhuang, Lizhen Huang.

**Data curation:** Shuiping Yan, Lizhen Huang, Xiaoqiang Xie, Lihong Chen.

**Formal analysis:** Shuiping Yan, Yimin Zhou.

**Funding acquisition:** Si Ai, Xiangli Zhuang.

**Investigation:** Shuiping Yan, Minfeng Yu, Lihong Chen.

**Methodology:** Shuiping Yan, Minfeng Yu, Xiangli Zhuang.

**Project administration:** Shuiping Yan, Jian Zheng, Si Ai, Xiangli Zhuang.

**Resources:** Shuiping Yan, Lizhen Huang, Xiaoqiang Xie.

**Supervision:** Shuiping Yan, Jian Zheng, Si Ai, Xiangli Zhuang, Minfeng Yu.

**Validation:** Shuiping Yan, Yimin Zhou, Xiangli Zhuang.

**Visualization:** Shuiping Yan, Xiangli Zhuang, Minfeng Yu.

**Writing – original draft:** Shuiping Yan.

**Writing – review & editing:** Shuiping Yan, Xiangli Zhuang, Minfeng Yu.

---

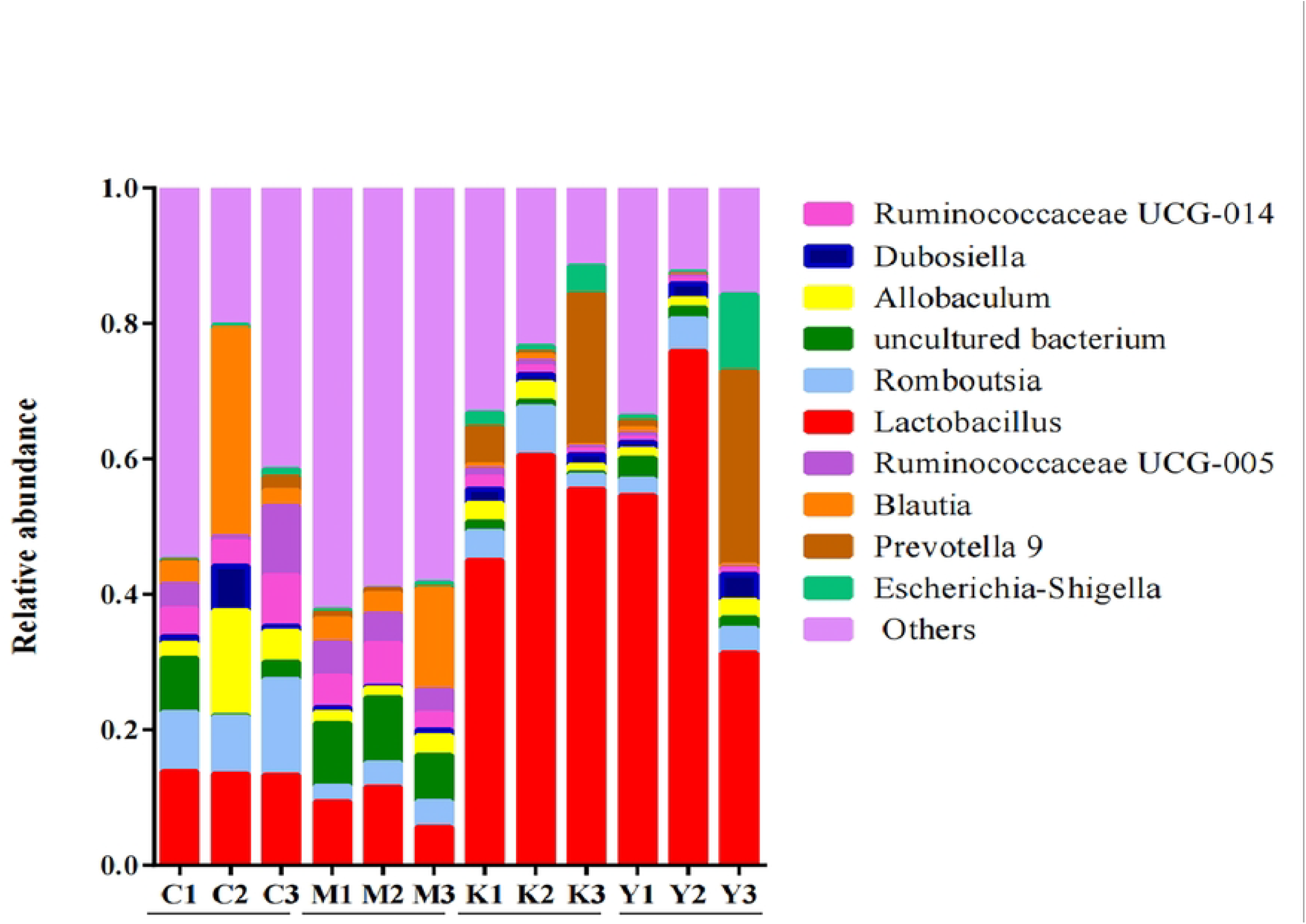

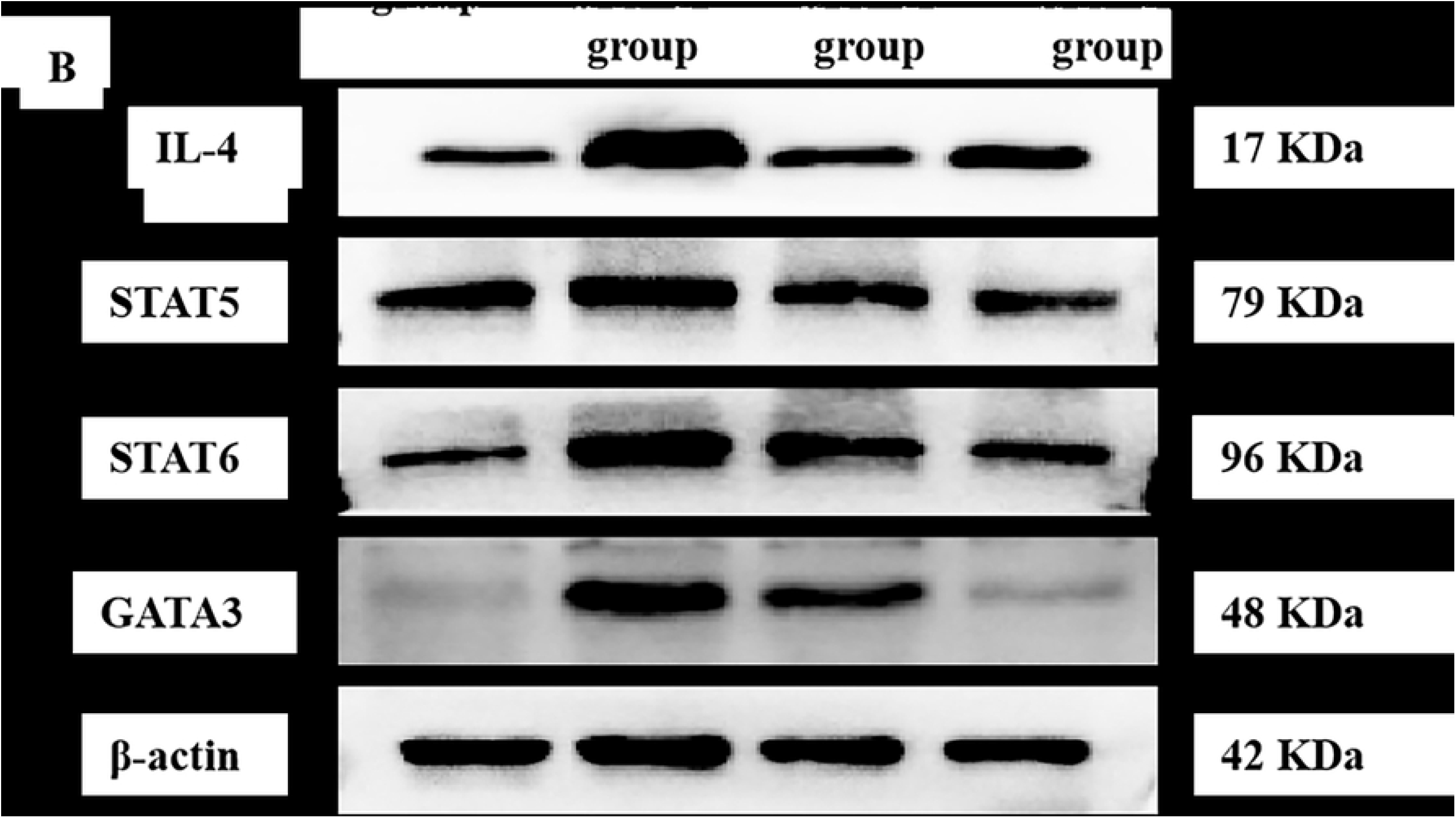

